# A large-scale bioinformatic study of graspimiditides and structural characterization of albusimiditide

**DOI:** 10.1101/2023.06.21.545981

**Authors:** Brian Choi, Arthur Acuna, Joseph D. Koos, A. James Link

## Abstract

Graspetides are a class of ribosomally synthesized and post-translationally modified peptides (RiPPs) that exhibits an impressive diversity in patterns of side chain-to-side chain ω-ester or ω-amide linkages. Recent studies have uncovered a significant portion of graspetides to contain an additional post-translational modification involving aspartimidylation catalyzed by an *O*-methyltransferase, predominantly found in the genomes of Actinomycetota. Here, we present a comprehensive bioinformatic analysis focused on graspetides harboring aspartimide for which we propose the name graspimiditides. From Protein BLAST results of 5,000 methyltransferase sequences, we identified 962 unique putative graspimiditides, which we further classified into eight main clusters based on sequence similarity along with several smaller clusters and singletons. The previously studied graspimiditides, fuscimiditide and amycolimiditide, are identified in this analysis; fuscimiditide is a singleton while amycolimiditide is in the fifth largest cluster. Cluster 1, by far the largest cluster, contains 641 members, encoded almost exclusively in the *Streptomyces* genus. To characterize an example of a graspimiditide in Cluster 1, we conducted experimental studies on the peptide from *Streptomyces albus* J1074, which we named albusimiditide. By tandem mass spectrometry, hydrazinolysis, and amino acid substitution experiments, we elucidated the structure of albusimiditide to be a large tetracyclic peptide with four ω-ester linkages generating a stem-loop structure with one aspartimide. The ester crosslinks form 22-, 46-, 22-, and 44-atom macrocycles, last of which, the loop, contains the enzymatically installed aspartimide. Further *in vitro* experiments revealed that the aspartimide hydrolyzes in a 3:1 ratio of isoaspartate to aspartate residues. Overall, this study offers a comprehensive insight into the diversity and structural features of graspimiditides, paving the way for future investigations of this unique class of natural product.

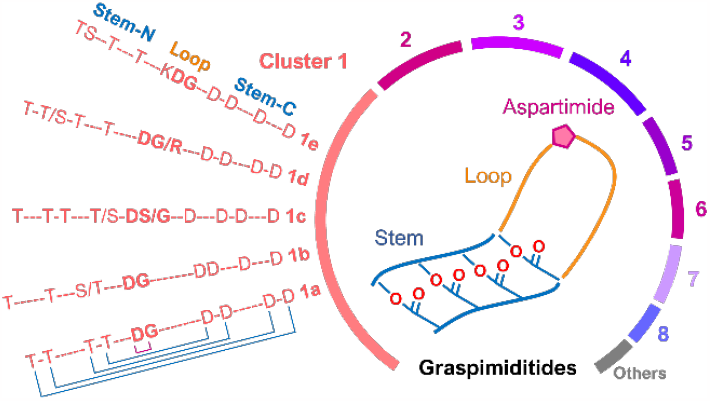

## Introduction

Natural products encompass a diverse array of chemical compounds that exhibit a wide range of biological activities.^1^ One such class of peptide-based natural products are ribosomally synthesized and post-translationally modified peptides (RiPPs). These peptides are derived from a polypeptide chain precursor which is genetically encoded with other biosynthetic genes. The N-terminal regions of precursor peptides, or leader peptides, often contain a motif that biosynthetic machinery recognizes to selectively modify the C-terminal portion, or core peptide. As this is a common mechanism employed by RiPP biosynthetic enzymes, the unique chemical moieties or connectivity they install on the C-terminal region of the precursor form the basis for classification. Nevertheless, considerable structural variability prevails within each class of RiPPs, calling for comprehensive, large-scale investigations to describe their diversity. Over the past decade, such studies have become increasingly feasible, owing to the development of high-throughput RiPP genome mining pipelines that can be applied across the rapidly expanding wealth of available genomic data.^2-6^

A class of RiPPs, known as graspetides, has recently expanded from large-scale genome mining studies since 2020.^7-9^ These studies revealed thousands of graspetide biosynthetic gene clusters (BGCs), with the highest number reported as 4,356 (3,455 unique) precursors from 3,923 BGCs identified by Ramesh, et al.^8^ Experimental work has characterized the chemical structures of several representative graspetides,^7-8, 10-14^ collectively showcasing an immense diversity in the patterns of side chain-to-side chain ω-ester or ω-amide crosslinks that form complex, macrocyclic structures (Figure 1A).^15^ Previously, knowledge of graspetides was limited to a subset called microviridins, which are serine protease inhibitors with a tricyclic caged structure formed by two ω-ester and one ω-amide crosslinks.^15-17^ While various bacteria across the Cyanobacteria, Pseudomonadota (syn. Proteobacteria), and Bacteroidota (syn. Bacteroidetes) phyla produce microviridins, they exhibit relatively limited diversity in amino acid sequence and the same crosslinking pattern. Similar graspetides, known as marinostatins, have a microviridin-like crosslinking pattern, but lack the ω-amide linkage.^18-20^

**Figure 1.**
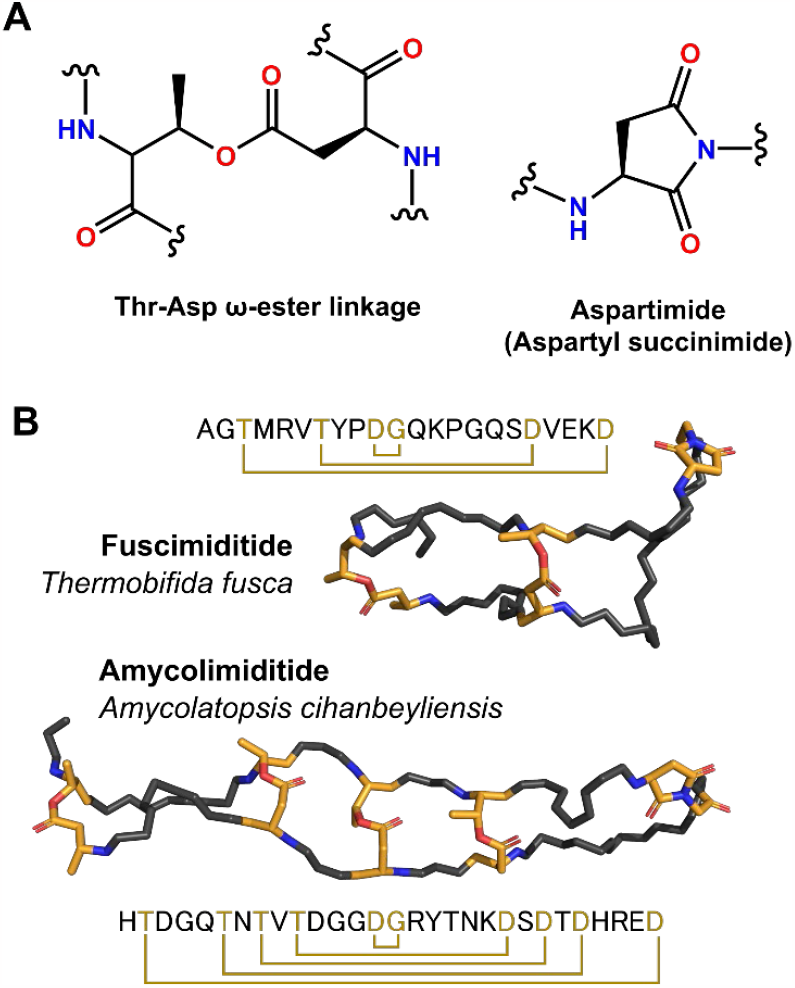
The structural basis of graspimiditides. (A) The chemical structure of (left) a Thr-Asp ω-ester linkage installed by an ATP-grasp enzyme and (right) an aspartimide installed by a RiPP-associated aspartyl *O*-methyltransferase. (B) NMR structures of the experimentally characterized graspimiditides, fuscimiditide (top, PDB: 7LIF) and amycolimiditide (bottom, PDB: 8DYM). The locations of ester linkages and aspartimides are indicated in the sequence maps.

Our research group recently discovered and characterized two graspetides, fuscimiditide and amycolimiditide, containing an additional post-translational modification: the cyclization of aspartate into aspartimide catalyzed by a methyltransferase (Figure 1A).^14, 21^ This moiety is located in the loop macrocycle of the stem-loop hairpin structure formed by sequence-wise parallel Thr-Asp ω-ester linkages (Figure 1B). Sequence-wise parallel here refers to the fact that the spacing of the residues involved in the ester crosslinks is equivalent on the N- and C-terminal stems. Two of the aforementioned large-scale genome mining studies, carried out by Ramesh et al. and Makarova et al., also identified and categorized a group of graspetide BGCs, a large portion of which contains such methyltransferases. These BGCs are predominantly found in the genomes of Actinomycetota, especially within the *Streptomyces* genus. Similarly, the BGCs of other classes of RiPPs, known as lanthipeptides and lasso peptides, that encode a methyltransferase for aspartimidylation are also predominantly found in Actinomycetota.^22-25^

In this study, we adopt a unique approach to large-scale graspetide genome mining, focusing on the methyltransferase gene to exclusively analyze putative graspetides with a predicted aspartimide moiety. Unlike previous graspetide genome mining studies, we exclude BGCs lacking a methyltransferase gene. To distinguish aspartimide-containing graspetides from those without it, we propose the name graspimiditide for this subset of **grasp**e**tides** that harbor an aspart**imid**e. Similarly, we propose the names lassimiditide and lanthimiditide for lasso peptides and lanthipeptides that harbor aspartimide moieties.^24-25^ We conduct large-scale genome mining on graspimiditides using an in-house methyltransferase-centric pipeline applied to a dataset of 5,000 graspimiditide-associated methyltransferases homologous to the gene found in *Streptomyces albus* J1074. The resulting ensemble of 962 unique graspimiditides are grouped into eight clusters based on sequence similarity. Our bioinformatic analysis demonstrates considerable diversity in the predicted patterns of ω-ester or ω-amide connectivity. In support of experimental studies on graspimiditides, we determined the connectivity of the graspimiditide encoded in the genome of *S. albus* J1074. Adopting the name of its producer, we refer to this peptide as albusimiditide. With bioinformatics insights, tandem mass spectrometry and hydrazinolysis experiments indicate a tetracyclic hairpin structure for albusimiditide, displaying both similarities and differences in structural features compared to fuscimiditide and amycolimiditide.

## Results and Discussion

### Bioinformatic study of graspimiditide biosynthetic gene clusters

For the bioinformatics study of graspimiditides, we focused our analysis on the biosynthetic gene clusters (BGCs) that contain three genes encoding the precursor, ATP-grasp enzyme, and *O*-methyltransferase. To ensure that the BGCs being analyzed are as exclusive as possible to graspimiditides, the amino acid sequence of the graspetide aspartyl *O*-methyltransferase in the *Streptomyces albus* J1074 strain (GsaM) was used as query in BlastP, instead of the conventional use of an ATP-grasp enzyme sequence as query for graspetide genome mining^7-9^. The search produced a list of 5,000 methyltransferase homologs (Supplemental Dataset 1), from which the genes unrelated to graspimiditide biosynthesis were further excluded out. This filtration step was done by examining the neighboring genes of each methyltransferase hit, testing their relevance to graspetide biosynthesis by searching for the key words “ATP-grasp” or “RiPP” in gene annotations. Then, we exerted another criterion that the filtered BGC contain three genes, presuming them to encode a precursor, an ATP-grasp enzyme, and a methyltransferase. The curated list contains 1,083 putative graspimiditide BGCs originated from the Actinomycetota phylum exclusively (Figure S1, Supplemental Dataset 2). 847 of them were found in the *Streptomyces* genus, comprising 78% of the identified graspimiditide BGCs. Our in-house BioPython workflow demonstrates similar performance to RODEO for mining graspimiditide BGCs, as the number of identified BGCs is close to the 1,326 BGCs identified as Group 13 graspetides by Ramesh et al^8^. Hence, while our genome mining approach relies on gene annotations, missing BGCs that are poorly described, the size of our curated list of graspimiditide BGCs is sufficiently large to conduct a large-scale comparison.

To construct the sequence similarity network (SSN) of the graspimiditide precursors, the amino acid sequences of precursors were extracted from the list of 1,083 graspimiditide BGCs and eliminated any redundancy with identical accession IDs, resulting in a list of 962 putative graspimiditide precursors. Only the last 50 residues were used for building the SSN, which grouped the collection into 8 major clusters, some small clusters and singletons (Figure 2A, Supplemental Dataset 3). Clusters 1, 3, and 4 were further divided into subclusters to facilitate the identification of sequence motifs for the core peptides, all of which are shown in Figure 2B, S1-S17. The previously studied graspimiditides, fuscimiditide (Figure 2C) and amycolimiditide (Figure 2D), is located as a singleton and in Cluster 5 of this SSN, respectively. The BGC containing the query methyltransferase from *S. albus* is found in Cluster 1a, a 49-membered subcluster of Cluster 1, the largest cluster comprised of 641 graspimiditides.

**Figure 2.**
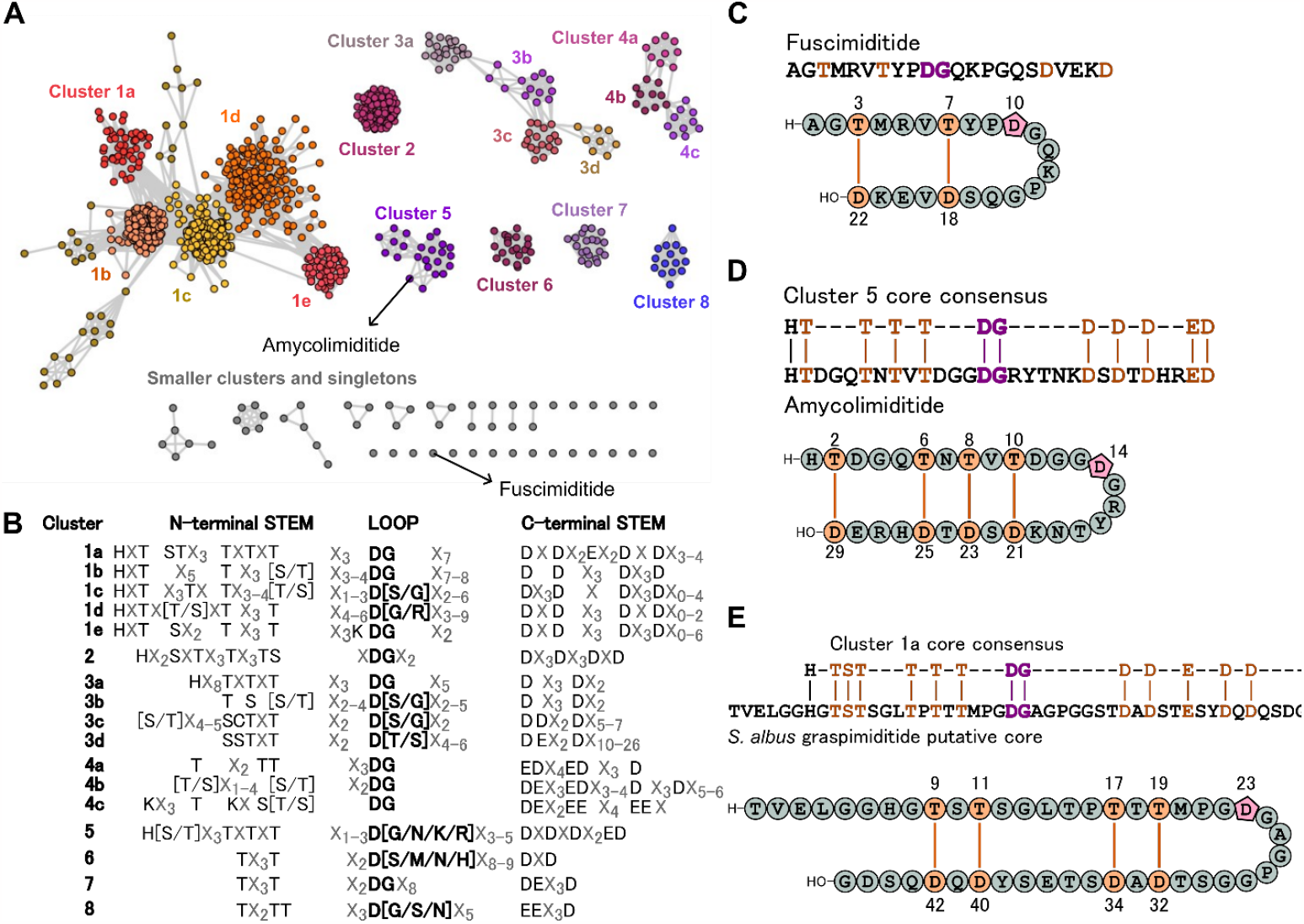
High throughput bioinformatics study of graspimiditides. (A) The sequence similarity network (SSN) of graspimiditide precursors, based on the last 50 amino acid residues. The ensemble is divided into 8 major clusters, some smaller clusters (≤ 6 members), and singletons. Amycolimiditide is found in Cluster 5 of the SSN, whereas fuscimiditide is isolated as a singleton. (B) The consensus sequence motifs of core peptides observed from each (sub)cluster, divided into three parts: N-terminal stem (left), loop (middle), and C-terminal stem (right). The N-terminal stem region is enriched with Thr and Ser residues, whereas the C-terminal stem region is enriched with Asp and Glu residues. In the loop region, the expected aspartimidylation site (i.e. the first conserved Asp-Xaa dipeptide) is bolded. Residues enclosed in square brackets represent co-occurring amino acids at a single position, listed in order of decreasing frequency. (C) The sequence of fuscimiditide with esterified residues colored orange and aspartimidylated residues colored purple. The cartoon beneath shows ester crosslinks in orange and the aspartimide represented as the pink pentagon. There are three residues between the ester crosslinks, both N- and C-terminally. (D) The sequence alignment of the Cluster 5 consensus motif and amycolimiditide with a cartoon representing amycolimiditide. Similar to fuscimiditide, amycolimiditide contains equal number of residues between all pairs of ester crosslinks. (E) The sequence alignment of the Cluster 1a consensus motif and the *S. albus* graspimiditide putative core. The cartoon represents a predicted ω-ester connectivity of the *S. albus* graspimiditide, based on the equal spacing of crosslinks observed in fuscimiditide and amycolimiditide.

The core peptide motifs of graspimiditides exhibit a wide range of sequence patterns. Like other classes of graspetides that do not contain the aspartimide, the core motif can be divided into an N-terminal region enriched in donor residues such as Ser, Thr, or Lys and C-terminal region enriched in acceptor residues Asp or Glu.^7^ However, the unique feature of graspimiditide core motif is the presence of intermediate region between the donor and acceptor-enriched regions, from which we identified the putative aspartimide sites. Strangely, the putative aspartimide site, which is the conserved Asp-Xaa dyad, appears to be globally located at or slightly N-terminal to the middle of this intermediate region. Previously, we had identified the *n*+1 aspartimide-follower residue (Xaa in Asp-Xaa) to be mostly Gly or occasionally Ser amongst a small subset of graspimiditides.^14, 21^ However, our bioinformatics study here shows unprecedented arginine, threonine, asparagine, lysine, methionine, and histidine *n*+1 residues, observed in Clusters 1d, 3d, 5, 6, and 8 (Figure 2B). While other residues in the intermediate region seem random, some graspimiditides are enriched with glycine in this region, such as GNPSDGSGPGGGGGG in WP_255929332.1 from *Streptomyces rubrisoli*. This suggests that amongst some graspimiditides, the flexibility of the intermediate region is important, either for biosynthesis or bioactivity. Another peculiarity was observed with Cluster 3d graspimiditides, which contain unusually long C-terminal tails. Additionally, graspimiditides in Clusters 1, 2, 3a, and 5 contain an exceptionally well-conserved hisitidine residue located upstream of the donor-enriched region in the core peptide motifs (Figure S2-S8, S15).

The graspimiditide precursor from *S. albus* J1074 is found in Cluster 1a, which exhibits the core consensus motif, HxTSTx_3_TxTxTx_3_DGx_7_DxDx_2_Ex_2_DxDx_3-4_ (Figure 2B, S1). In the alignment of the *S. albus* graspimiditide C-terminal sequence with the Cluster 1a core consensus motif, some residues can be predicted as esterified or aspartimidylated residues (Figure 2E). Designating the trypsin cut site as residue number 1, Thr9, Thr11, Thr17, and Thr19 are likely ω-ester donor residues, and Asp32, Asp34, Asp40, and Asp42 are likely ω-ester acceptor residues if the peptide follows the sequence-wise parallel esterification observed in other graspimiditides. The conserved Asp-Gly dipeptide located between the Thr/Ser- and Asp/Glu-enriched regions is the most likely aspartimidylation site. The implied geometry of the *S. albus* graspimiditide is two large macrocycles, each reinforced by two ω-ester linkages making a smaller macrocycle (Figure 2E). The sizes of the loop and stem macrocycles and the overall core peptide are larger than any graspetide core previously characterized. Additionally, we expect some structural complexity from the presence of many proline residues, one located in the putative stem region and two located in the putative loop. The presence of nine potential Ser or Thr residues in the N-terminal stem merits further experimental investigation in determining the residues used for the construction of ω-ester crosslinks in the mature graspimiditide product. This would offer further insight into the substrate specificity of graspetide ATP-grasp enzymes, an on-going topic being investigated in RiPPs research^16, 26-27^.

### Reconstitution of *S. albus* J1074 graspimiditide biosynthesis

Based on the expected structural novelty and widespread existence of similar peptides, the *S. albus* J1074 graspimiditide, which we name albusimiditide, was pursued further for experimental characterization. We used *E. coli* to coexpress the precursor gene with the graspimiditide maturation enzymes (ATP-grasp enzyme and *O*-methyltransferase) to assess the extent of modifications. When coexpressed with the ATP-grasp enzyme GsaB, the precursor protein GsaA exhibited an average mass of 10062.3 Da (calculated theoretical average mass = 10062.7 Da), a -72 Da shift from the expected unmodified mass corresponding to 4-fold dehydration (Figure 3A). Coexpressing GsaA with GsaB and the methyltransferase GsaM resulted in an observed average mass of 10044.5 Da (calculated theoretical average mass = 10044.6 Da) from the modified precursor, exhibiting a -90 Da shift corresponding to 5-fold dehydration (Figure 3A). To verify no background post-translational modification in the *E. coli* host used, the precursor protein GsaA was expressed alone, and the expected unmodified average mass of 10134.3 Da was observed (calculated theoretical average mass = 10134.7 Da). Overall, the coexpression studies suggest that GsaB installs four crosslinks and GsaM catalyzes one cyclic imide formation.

**Figure 3.**
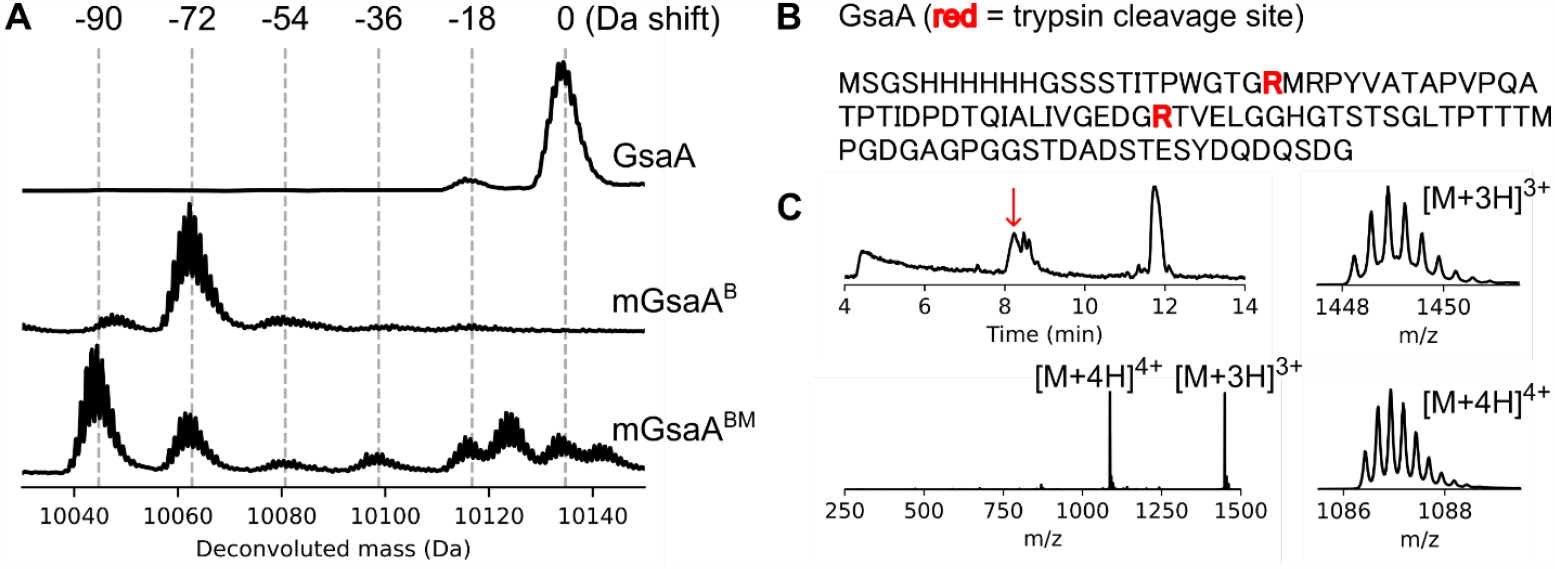
Enzymatic modifications to the albusimiditide precursor. (A) Coexpression studies of the precursor protein with graspimiditide maturases. The deconvoluted mass spectra of the (un)modified precursor protein from (co)expression is shown. (top) GsaA, the precursor protein expressed alone. (middle) mGsaA^B^, GsaA modified by the ATP-grasp enzyme GsaB, exhibiting four dehydrations. (bottom) mGsaA^BM^, GsaA modified by GsaB and RiPP aspartyl *O*-methyltransferase GsaM, exhibiting five dehydrations. (B) Identification of trypsin cleavage sites (red) in the GsaA sequence. (C) Trypsin digestion of mGsaA^B^ for core peptide isolation. (top left) The total ion chromatogram (TIC) of the mGsaA^B^ trypsin digestion products. The red arrow indicates the 8.2 min peak corresponding to the four-fold dehydrated 46 aa C-terminal peptide (pre-albusimiditide). (bottom left) Mass spectrum extracted across 8.1-8.3 min showing +3 and +4 charged species of pre-albusimiditide. (right) Isotopic distributions of the (top) +3 and (bottom) +4 charged species.

To identify and isolate the core peptide of albusimiditide, we used trypsin to digest the leader peptide of mGsaA^B^ (GsaB-modified GsaA). The amino acid sequence of GsaA contains only two possible trypsin cleavage sites, which are Lys or Arg residues followed by any residue other than proline (Figure 3B). Analyzing the digested products by LC-HRMS, we identified a peak chromatographed at 8.2 min retention time containing a +3 charged species with m/z value of 1448.24 (expected value: 1448.26) and a +4 charged species with m/z value of 1086.43 (expected value: 1086.44), corresponding to the four-fold dehydrated 46 aa long C-terminal peptide of GsaA (Figure 3C). This peptide was isolated to a high purity by HPLC (Figure S19). To further trim the N-terminal end of the tryptic peptide, the HPLC-purified product was lyophilized and further digested with aminopeptidase. This reaction digested away four N-terminal residues Thr-Val-Glu-Leu, verified by the +3 charged species with m/z value of 1300.85 (expected value: 1300.85) and a +4 charged species with m/z value of 975.89 (expected value: 975.88) observed by mass spectrometry (Figure S20). Performing the aminopeptidase digestion longer did not tailor the peptide further.

Based on the presence of the Gly-Gly dipeptide sequence in the remaining peptide, we expected the LahT150 peptidase domain of AMS/PCAT transporter, commonly employed exogenously in RiPP characterization studies^28-30^, could recognize the GG motif in mGsaA^B^ and cleave before the conserved His residue (Figure 3B). The protease, however, did not process mGsaA^B^, leaving the crosslinked protein intact. Then, we turned to trying endogenous proteases found in the same producer *S. albus* J1074. We mixed the purified mGsaA^B^ protein with the supernatant of the spent liquid growth medium of *S. albus* J1074. From the LC-HRMS analysis, we detected a peak with identical mass corresponding to the quadruply dehydrated 42 aa C-terminal peptide observed in the aminopeptidase experiment (Figure S21). From the leader peptide proteolysis experiments, we infer that the 42 aa core peptide is the intended size of albusimiditide. Further structural analysis of the graspimiditide was pursued with the 46 aa tryptic core peptide for experimental simplicity. In this study, we name the tryptic core peptide with the four ester crosslinks pre-albusimiditide and a similar peptide with the GsaM-installed aspartimide albusimiditide. We note that this naming convention is used in all imiditide studies from our research group^14, 21, 25^.

### Mapping the ω-ester crosslinks installed by GsaB

Finding the locations of all PTMs installed by maturase enzymes is a major task towards solving the chemical structure of a RiPP natural product. For graspetides, the ^1^H-^13^C HMBC is the golden standard experiment for mapping the ω-ester or ω-amide linkages installed by ATP-grasp enzymes.^14-15, 21, 31^ However, we expected the 2D NMR approaches we and others have previously used^14, 21, 31^ to be not suitable for albusimiditide. At 46 aa, albusimiditide is more than 50% longer than amycolimiditide (Figure 2), which is the largest graspimiditide analyzed by 2D NMR experiments so far. Moreover, it harbors multiple Gly, Ser, Thr, and Pro residues. Especially with three proline residues, the isolated peptide could contain different combinations of *cis* or *trans* proline isomers, further complicating the chemical shift assignments. Therefore, we turned to an iterative approach based on mass spectrometry, hydrazide labeling, and mutagenesis to map the locations of all four crosslinks (Figure 4).

**Figure 4.**
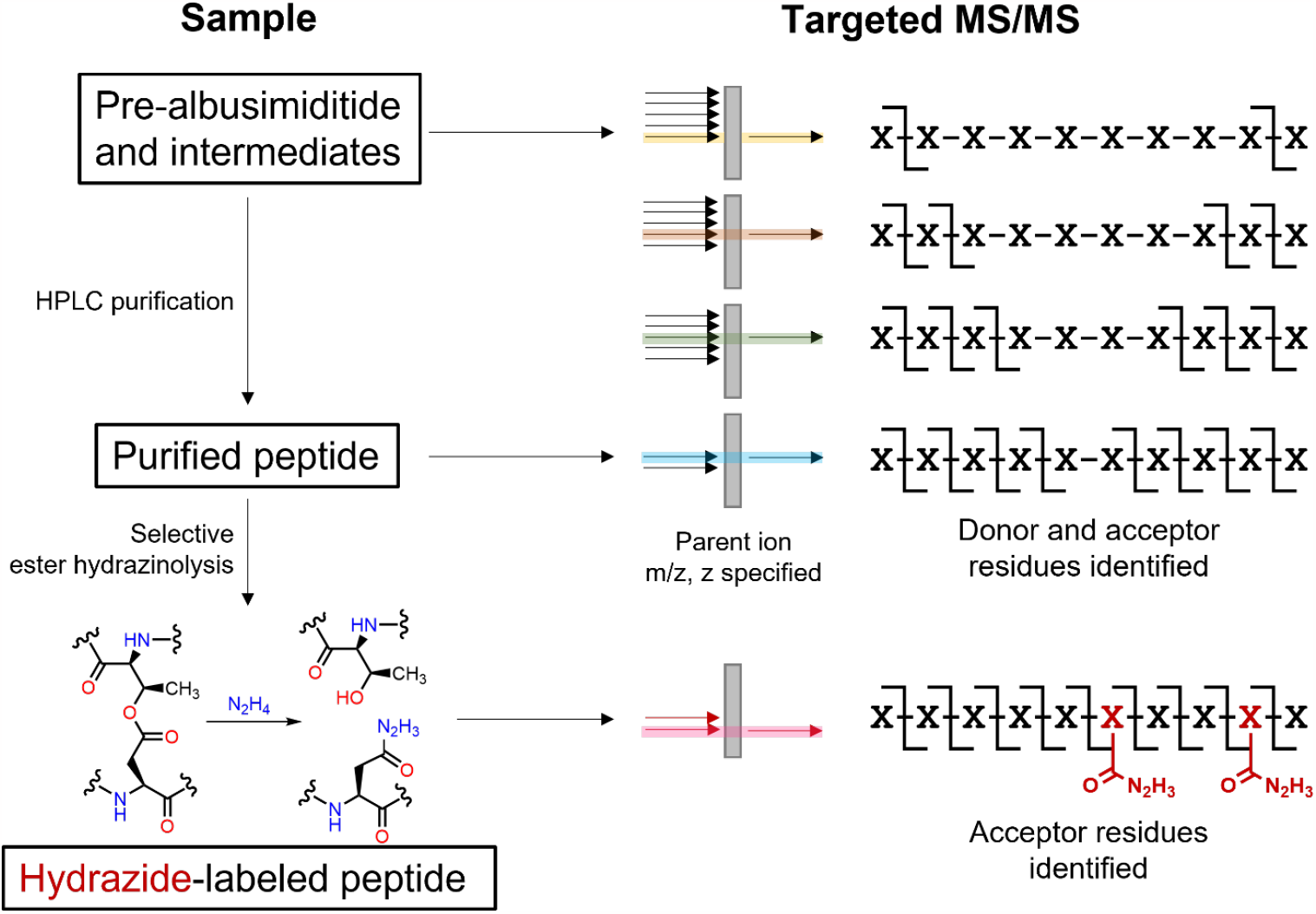
Strategy for mapping the ω-ester linkages of albusimiditide. Collision-induced dissociation (CID) is targeted on a parent ion with specified m/z and z values. The sample could be a crude mixture of mGsaA^B^ trypsinate or an (ester-hydrazinolyzed) HPLC-purified core peptide. The experiments are iterated over pre-albusimiditide and its intermediates, and the MS/MS spectra are analyzed comparatively to assign the ω-ester linkages, based on emerging daughter ions and hydrazide-labeling.

In this strategy, collision induced dissociation (CID) is performed on pre-albusimiditide and partially esterified intermediates. We compare the MS/MS fragmentation patterns of the different intermediates and search for the emerging MS/MS fragments to assign the ω-ester linkages. In the tandem mass spectrometry experiments, a precursor ion is selected with specified m/z and z values, obviating the need to purify the core peptides as single species. We purify those that can be isolated in high purity, such as pre-albusimiditide (Figure S19), to obtain a high-quality MS/MS spectrum. We additionally perform selective ester hydrazinolysis followed by CID on the HPLC-purified peptides for further verification of crosslinked acceptor residues.

To scope out where the donor and acceptor residues for the four ω-ester linkages are localized, the CID experiment was performed on pre-albusimiditide. From the resulting high-quality MS/MS spectrum, we observed b-ion fragments resulting from concomitant N-terminal fragmentation at Pro21 (which we name b** ions) and Pro27 (b***), and not Pro16 (b*) (Figure 5A, S22). This is an acknowledged phenomenon known as the “proline effect”, frequently observed in CID experiments performed with an electrospray ionization mass spectrometer (ESI-MS) (Figure S23)^32-34^. The fragmentation pattern reveals two regions where MS/MS fragments are not observed: N-terminal region Thr9-Thr19 enriched in Thr and Ser residues and C-terminal region Ala33-Asp42 enriched in Asp and Glu residues. Four pairs of donor and acceptor residues for ω-ester linkages are most likely to be found in these regions. When all four crosslinks are hydrazinolyzed, the CID experiment yields significantly more MS/MS fragments. Importantly, b* ions (resulting from the proline effect at Pro16) are detected, unlike the MS/MS experiment of pre-albusimiditide with all ester linkages intact (Figure 5A). The revelation of b* ions is consistent with our hypothesis that the four ester linkages are localized between Thr9 and Thr19. Various mass shifts in the daughter ions demonstrate the following pattern: +14 Da shift was observed in all fragments containing Asp32, +28 Da in all ions containing Asp32 and Asp34, +42 Da in all ions containing Asp32, Asp34, and Asp40, and +56 Da in all ions containing Asp32, Asp34, Asp40, and Asp42 (Figure 5A, S24). All ions not containing Asp32 exhibited unmodified m/z values. The pattern supported by several MS/MS fragments clearly indicate that Asp32, Asp34, Asp40, and Asp42 are the acceptor residues of the four ω-ester linkages.

**Figure 5.**
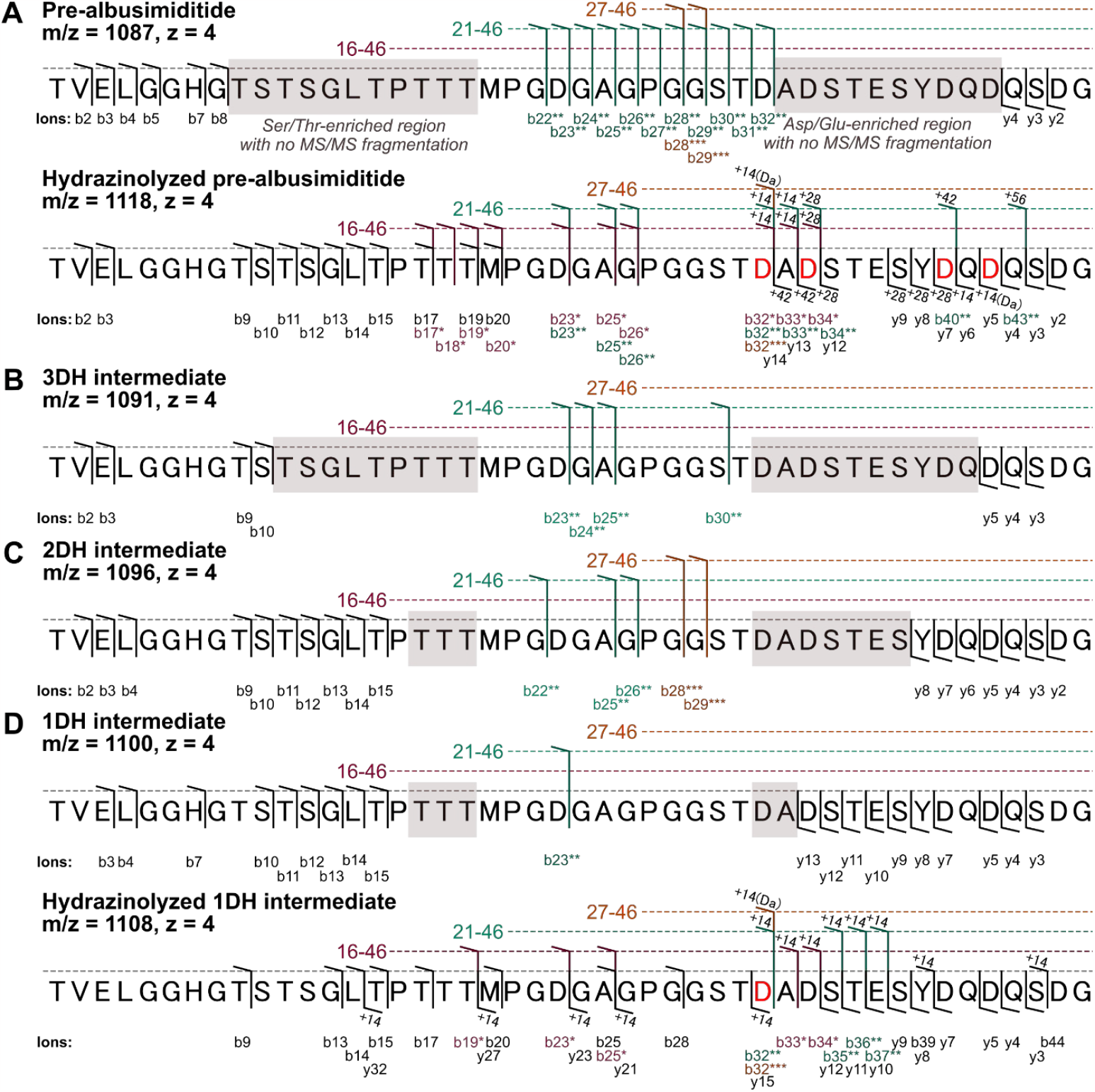
Mapping the ω-ester crosslinks in albusimiditide. MS/MS fragmentation of (A, top) pre-albusimiditide and (bottom) its hydrazinolysate, and the core peptides of pre-albusimiditide intermediates harboring (B) 3 dehydrations (3DH), (C) 2 dehydrations (2DH), and (D, top) 1 dehydration (1DH) and (bottom) its hydrazinolysate. For panels with two fragmentation diagrams, the bottom one corresponds to the ester hydrazinolysis experiment, and the mass shift values associated with the presence of hydrazide adduct(s) are indicated at the fragmentation markers. The implicated esterified acceptor residues are colored red. The colors and special b-ion nomenclature, magenta and b*, teal and b**, and gold and b***, are used for fragments associated with the proline effect at Pro16, Pro21, and Pro27, respectively. The no-fragmentation regions enriched in candidate donor (Ser/Thr) and acceptor residues (Asp/Glu) for ω-ester linkages are shaded grey.

On a minor note, the b32** ion detected in the CID experiment of pre-albusimiditide obtained from the C-terminal fragmentation of the esterified Asp32 residue, is detected. This ion could emerge from a potential McLafferty rearrangement that undoes the ω-ester linkage containing Asp32. This type of ester rearrangement is a common occurrence in MALDI-TOF-MS/MS analysis of graspetides with ω-ester linkages, leading to a net loss of H_2_O from esterified Ser or Thr residues^7, 10^-^11, 30^. However, in our MS/MS analysis, we intentionally disregard this -18 Da shift effect when identifying and assigning ω-ester donor residues. This is because albusimiditide contains numerous Ser and Thr residues, which are widely recognized to undergo dehydration when present as free residues in tandem mass spectrometry^35-37^.

Then, we performed CID on the 3DH intermediate core peptide to search for b and y ions not observed in the pre-albusimiditide CID experiment. The MS/MS spectrum revealed b_9_, b_10_, and y_5_ ions, which are not detected from the pre-albusimiditide MS/MS experiment (Figure 5B, S25). We reason that Thr9 and Asp42 form the ω-ester linkage lacking in the 3DH intermediate. To verify this assignment, we coexpressed GsaA harboring T9V and D42N substitutions with GsaB. The precursor protein was dehydrated three times maximally, consistent with the Thr9-Asp42 linkage assignment (Figure S26).

To identify the second ω-ester linkage, CID was performed on the 2DH intermediate core peptide. The EIC of the 2DH intermediate showed two separate peaks (Figure S27), and the MS/MS spectrum corresponding to the major product (8.1-8.2 min retention time in LC) was extracted for analysis. The spectrum displayed a more widespread fragmentation across the albusimiditide sequence than observed in the MS/MS spectrum of pre-albusimiditide or the 3DH intermediate (Figure 5C, S28). The observed C-terminal fragmentation past Thr11 (b_11_, b_12_, b_13_, b_14_, and b_15_) and N-terminal fragmentation past Asp40 (y_6_, y_7_, and y_8_), not detected in the CID experiment of the 3DH intermediate, suggests a Thr11-Asp40 ω-ester crosslink. The extended N and C-terminal fragmentation is blocked at Pro16 and Ser38, leaving Thr17, Thr18, and Thr19 as candidate donor residues for the two-remaining ω-ester linkages involving Asp32 and Asp34. While Ser35, Thr36, and Ser38 cannot be strictly ruled out, all microviridins and other graspetides whose crosslinks were located experimentally collectively demonstrate a pattern, in which a donor residue must be located N-terminal to its paired acceptor residue^7^. To verify the Thr11-Asp40 linkage, we coexpressed GsaA harboring T11V and D40N substitutions with GsaB. The mGsaA^B^ variant exhibited two dehydrations (Figure S29). This is similarly observed with the amycolimiditide variant whose second outermost ester linkage is substituted with isosteric amino acid residues^21^. The additional loss of dehydration occurs because the formation of the outermost ω-ester linkage depends on the presence of the second outermost crosslink. To test whether this is also true for albusimiditide, the core peptide of the T11V D40N variant was further analyzed by tandem mass spectrometry after trypsin digestion and subsequent HPLC purification (Figure S30). CID yielded an MS/MS spectrum showing fragmentation past the region containing the outermost crosslink, similar to the MS/MS spectrum of the 2DH pre-albusimiditide intermediate (Figure S31). We also point out that the y_9_, y_10_, and y_12_ ions are detected for this variant, ruling out Ser35, Thr36, and Ser38 as ω-ester donor residues for the two innermost ester linkages as expected.

To identify the two-remaining ω-ester linkages, we performed CID on the 1DH intermediate purified by HPLC (Figure S32). The resulting MS/MS spectrum indicated the presence of y_13_ ion not observed in the CID experiment of the 2DH intermediate (Figure 5D, S33). This indicates that the second ω-ester linkage formed on albusimiditide involves Asp34. However, no new b ion that would facilitate the assignment of the donor residue for the second linkage was detected. We sought a different approach by using elastase to digest the linear ends of the 1DH peptide. This reaction yielded a product with a mass of 2799.96 Da (expected mass: 2800.04 Da), corresponding to the 1DH pre-albusimiditide intermediate cleaved after Thr17 (Figure S34). This indicates that Thr17 is connected to Asp34. Since threonine is not known to be recognized by elastase, we sought to further verify the identity of the 2800 Da molecule by MS/MS. The resulting spectrum showed b** and y ions consistent with the amino acid sequence of albusimiditide (Figure S35).

Thr18, Thr19, and Asp32 are the remaining residues to be selected for assigning the last remaining ω-ester linkage, which is the first crosslink installed on albusimiditide. To verify Asp32 as the acceptor residue for this linkage, ester hydrazinolysis was performed on the 1DH intermediate, followed by CID. In the MS/MS spectrum, all daughter ions containing Asp32 exhibited +14 Da upshift, verifying Asp32 as the acceptor residue for the first ω-ester linkage of albusimiditide (Figure 5D, S36). Since we could not conclusively determine the donor residue of the first ω-ester linkage by MS/MS, we attempted to solve this by mutagenesis approach. Coexpressing each of the two GsaA variants, T18V D32N and T19V D32N, with GsaB resulted in single and no dehydration, respectively (Figure S37, S38). Since mutagenesis of the first crosslink installed by an ATP-grasp enzyme commonly ablates the formation of all other crosslinks among graspetides,^21, 38^ we assigned Thr19-Asp32 to be the first ω-ester linkage of albusimiditide. For additional verification, we tested another variant with T18V substitution targeting the putative noncrosslinking residue. The coexpression experiment showed mostly four-fold dehydration, exhibiting no disruption in esterification as expected (Figure S39).

### Characterization of the aspartimide in albusimiditide

With high confidence, we have mapped the four ω-ester linkages of albusimiditide to be T19-D32, T17-D34, T11-D40, and T9-D42, indicating a tetracyclic hairpin structure like amycolimiditide but with a 44-membered loop, one large, 46-membered stem macrocycle and two small, 22-membered stem macrocycles (Figure 6A). As a final step towards determining the structure of albusimiditide, we sought to identify the location of aspartimide installed by GsaM. The sequence alignment of precursors in Cluster 1A graspimiditides shows Asp23-Gly24 to be exceptionally well-conserved (Figure S2). We expected this conserved dipeptide to be the aspartimidylation site, similar to the DG dipeptides that are aspartimidylated in fuscimiditide and amycolimiditide^14, 21^. For experimental verification, mGsaA^B^ and mGsaA^BM^ for the D23N variant were analyzed by mass spectrometry, which indicated four dehydrations observed from both proteins (Figure 6B). The lack of aspartimidylation indicates that Asp23 is the designated aspartimide location for albusimiditide. We incorporated this assignment in the solved chemical structure of albusimiditide, shown in Figure 6A.

**Figure 6.**
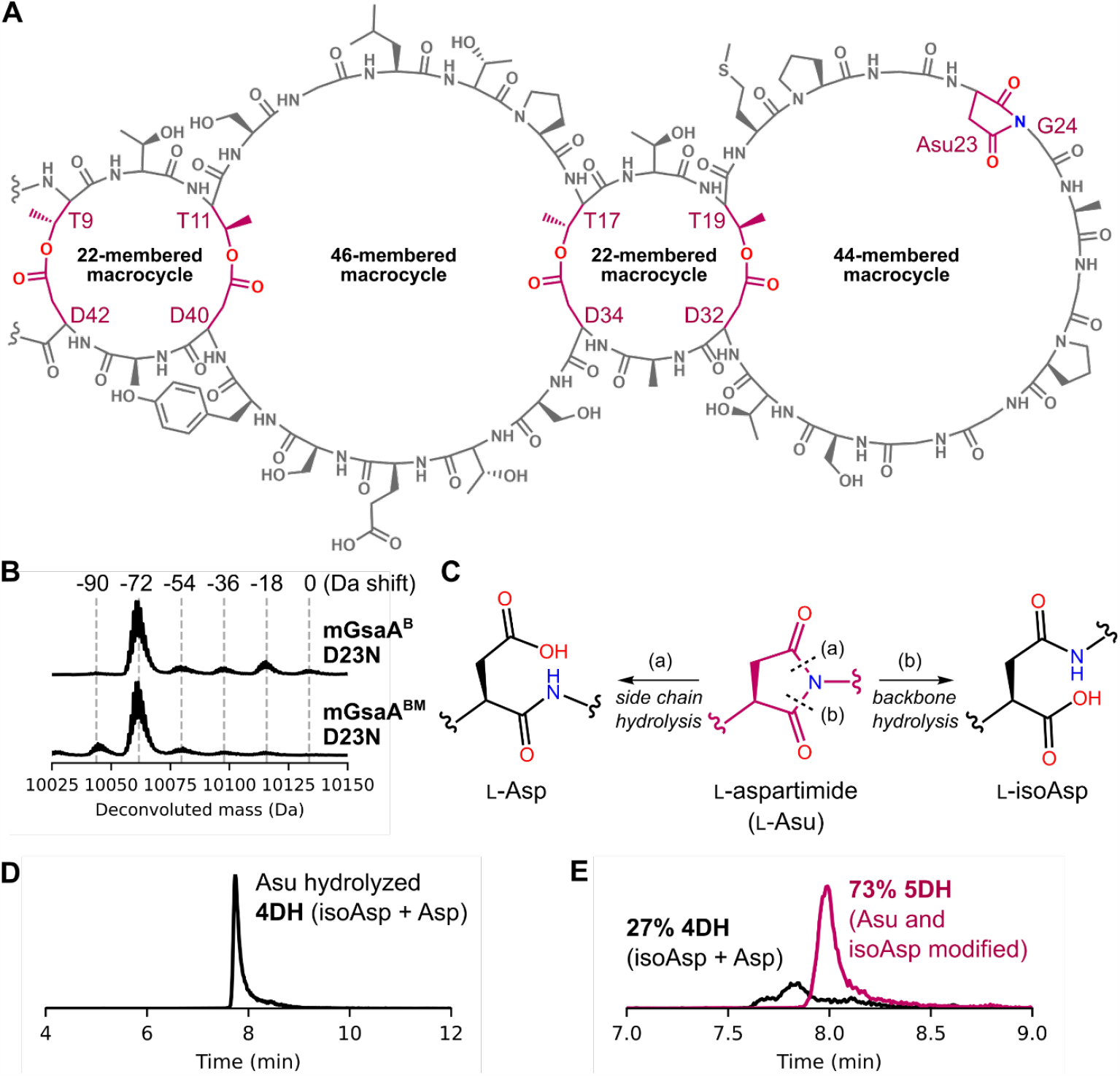
The characteristics of albusimiditide. (A) The experimentally solved chemical structure of the macrocyclic portion of albusimiditide. The ω-ester linkages and the aspartimide are colored maroon. The linkages are T19-D32, T17-D34, T11-D40, and T9-D42. The aspartimide is the 23^rd^ residue located in the loop macrocycle. (B) The comparison of deconvoluted mass spectra of (top) mGsaA^B^ and mGsaA^BM^ harboring the D23N substitution. The nearly identical spectra showing four-fold dehydrated species as the major product implies Asp23 to be aspartimidylated by GsaM. (C) Chemical pathway for aspartimide hydrolysis resulting in the formation of Asp and isoAsp. (D) Liquid chromatography of albusimiditide with aspartimide hydrolyzed, showing a single peak for the two expected hydrolysate products. (E) Extracted ion chromatograms for 4DH and 5DH albusimiditide species, after in vitro reaction of aspartimide-hydrolyzed albusimiditide species with *H. sapiens* protein L-isoaspartyl *O*-methyltransferase (*Hs*PIMT). The regioselectivity of hydrolysis is 73% towards isoAsp, based on the EIC peak area.

Then, we sought to characterize the propensity of isoaspartate (isoAsp) formation resulting from the aspartimide hydrolysis at the peptide backbone (Figure 6C). The regioselectivity of aspartimide hydrolysis (i.e. the ratio of isoAsp vs Asp formation upon aspartimide hydrolysis) is often a characteristic feature idiosyncratic to individual RiPP imiditides. For example, the aspartimide in fuscimiditide hydrolyzes mostly to isoAsp at 95% regioselectivity^14^, whereas the aspartimide in the lassimiditide lihuanodin exclusively hydrolyzes to aspartate^25^. This complete reversion of aspartimide to aspartate has further structural implication for lihuanodin, as a mechanism for maintaining its threaded rotaxane structure^39^. In general, RiPP peptides with hydrolyzed aspartimide tend to chromatograph as two distinct peaks corresponding to aspartimide hydrolyzing to isoAsp or Asp. For albusimiditide, however, we observed one distinct peak with retention time corresponding to pre-albusimiditide after aspartimide hydrolysis (Figure 6D). To consider the possibility that isopre-albusimiditide (albusimiditide with isoAsp at the aspartimide site) chromatographed similarly to pre-albusimiditide, we performed an in vitro reaction with *H. sapiens* PIMT to revert any isoAsp present to aspartimide. After the reaction, approximately 73% of the product was aspartimidylated (Figure 6E), exhibiting a typical ∼7:3 isoAsp:Asp regioselectivity of aspartimide hydrolysis observed from linear peptides. This regioselectivity is similarly observed with the lanthimiditide OlvA(BCS_A_) and the graspimiditide amycolimiditide^21, 24^.

## Conclusions

Here, we have conducted a comprehensive bioinformatic analysis of graspimiditides, which are a group of graspetides that contain an aspartimide moiety enzymatically installed by a dedicated L-aspartyl *O*-methyltransferase. Our genome mining workflow identified a diverse group of nearly thousand graspimiditides, all encoded in actinobacterial genomes (Figure S1). This specificity of origin could imply that their bioactivities are associated with the unique biological features of actinobacteria, including sporulation, mycelial growth, plant-microbe interaction, and high GC content.^40^ The bioactivities of RiPPs containing aspartimides have long been postulated to possess a single intended bioactive form, derived from one of three structural isoforms: aspartate, aspartimide, or isoaspartate.^14-15, 21, 24-25^ One could predict the isoAsp isoform to be the bioactive form, as most aspartimides in RiPPs and synthetic peptides exhibit regioselective aspartimide hydrolysis favoring isoAsp formation.^21, 24, 41-42^ Our findings with albusimiditide support this hypothesis, as *in vitro* selective isoAsp aspartimidylation experiments with *Hs*PIMT showed a roughly 3:1 ratio of isoAsp to Asp (Figure 6E). However, the aspartimide in the lasso peptide lihuanodin exclusively hydrolyzes to aspartate, in stark contrast to other experimentally characterized RiPPs containing aspartimides.^25^ A recent study by Cao et al. supports the notion that the reversion of aspartimide only to aspartate is intended to maintain the lasso peptide’s class-defining threaded lasso structure.^39^

Characterizing graspetides and graspimiditides involves identifying the locations of ω-ester and ω-amide linkages. For albusimiditide, we employed an extensive combination of mass spectrometry and hydrazinolysis experiments to determine its ω-ester crosslinking pattern (Figure 5), with equal numbers of N- and C-terminal interstitial residues between crosslinks (Figure 6A). This pattern is also observed in fuscimiditide and amycolimiditide, found to be only distantly related to albusimiditide based on the sequence similarity network (Figure 2A). Notably, unlike fuscimiditide and amycolimiditide, one of the stem macrocycles in albusimiditide is larger than the loop macrocycle (Figure 6A). Albusimiditide’s tetracyclic hairpin structure was predicted *a priori* by pairing the most conserved donor and acceptor residues in a manner that maintains equal numbers of interstitial residues on both N- and C-terminal sides (Figure 2C). Supported by our findings, we hypothesize that this method for predicting the mapping of ω-ester and ω-amide crosslinks may be broadly applicable to graspimiditides. We believe this would facilitate future graspimiditide characterization, particularly given their high serine and threonine content, some of which do not form ω-ester linkages. This is supported by the fact that many serine and threonine residues are not as well-conserved as the predicted ω-ester donor residues (Figures S2-S18).

Typically, RiPPs undergo further processing, including leader peptide excision and post-biosynthesis export.^3, 43^ Amongst graspetides, microviridins are the only group experimentally shown to have dedicated genes in their BGCs for leader peptide removal and export. The BGCs of other groups of graspetides contain biosynthetic genes only^12, 14, 21^ or harbor putative accessory genes awaiting experimental characterization.^8, 10^-^11, 13^ In the case of albusimiditide, we found that the supernatant of a culture of the putative producing strain can digest the leader peptide (Figure S21). Although this result does not provide definitive evidence for a non-microviridin-type graspetide-specific mechanism for leader peptide removal and export, it may suggest that graspetide BGCs have evolved to lose those accessory genes, relying instead on host-native proteases and transporters. Many questions regarding the origin, functions, and purposes of graspimiditides remain unanswered, calling for further experimental investigations beyond characterizing peptide structures.

## Methods

Detailed information on materials and methods is provided in Supporting Information File 1.

### Plasmid construction

The lists of oligonucleotides, plasmids generated, and cloning methodologies used in this study are provided in Tables S1-S3, respectively.

### Graspimiditide precursor protein production in *E. coli* and purification

The *E. coli* BL21 DE3 Δ*slyD* strain was electroporated with a plasmid harboring the graspimiditide precursor gene *gsaA* (wild-type or variant) and, optionally, *gsaB* (ATP-grasp enzyme) or *gsaBM* (ATP-grasp enzyme and *O*-methyltransferase). Transformants were grown on LB agar medium (5 g L^-1^ yeast extract, 10 g L^-1^ tryptone, 10 g L^-1^ NaCl, 15 g L^-1^ agar) supplemented with 100 μg mL^-1^ ampicillin (amp). A single colony was used to inoculate the 5 mL starter liquid LB+amp medium and then grown with shaking at 37 °C overnight. The starter culture was then back-diluted in 500 mL LB+amp medium to 0.02 OD_600_ and grown with shaking at 37 °C. Upon reaching 0.5 OD_600_, 1 mM isopropyl β-D-thiogalactoside (IPTG) was added to induce protein production. The culture was incubated at room temperature with shaking for two days and then centrifuged at 4,000 x *g* for 15 min. The cell pellet was resuspended with 10 mL urea buffer (8 M urea, 100 mM NaH_2_PO_4_, 10 mM Tris-base), pH 8.0, frozen at -80 °C, and then thawed at room temperature. The cell lysate was clarified by centrifugation and incubated with 1 mL Ni-NTA resin (Qiagen). The precursor protein was purified using the standard protocol from Qiagen for purifying protein from Ni-NTA resin under denaturing conditions with urea. For a long-term storage, the protein was buffer-exchanged in 1X PBS buffer (137 mM NaCl, 2.7 mM KCl, 10 mM Na_2_HPO_4_, 1.8 mM KH_2_PO_4_) and concentrated down with a 10 kDa cut-off centrifugal filter (Millipore).

### Graspimiditide core peptide purification

GsaB-modified GsaA precursor protein was digested with trypsin (Promega, sequencing-grade) at 37 °C for 20 min., following the supplier’s protocol. The reaction was quenched by adding 0.1 volume of 10% formic acid. The trypsin digestion mixture was subjected to HPLC (Agilent 1200 series), and the peak(s) corresponding to the desired core peptide product was collected. The core peptide was lyophilized and reconstituted in ultrapure water for subsequent experiments.

### Albusimiditide ω-ester connectivity analysis

The connectivity of ω-ester linkages in albusimiditide was determined by tandem mass spectrometry experiments on pre-albusimiditide and intermediates. Mass spectra were obtained with Agilent 1260 Infinity II LC and Agilent 6530 QTOF, with the CID voltage set by the formula V = 0.036 × (m/z) – 4.8. The MS2 spectrum of pre-albusimiditide was acquired from the HPLC-purified pre-albusimiditide sample obtained from the trypsin digest of mGsaA^B^. The MS2 spectrum of ester-hydrazinolyzed pre-albusimiditide was obtained from the crude sample of ester-hydrazinolyzed pre-albusimiditide, prepared by mixing equal volume of HPLC-purified pre-albusimiditide with the hydrazinolysis buffer (60% 1X PBS, pH 5.0, 40% 35 wt% hydrazine), pH ∼8 and incubating at 55 °C for 45 min. The MS2 spectrum of the 3DH and 2DH intermediates of pre-albusimiditide was obtained from the trypsin digest of mGsaA^B^, as low amount species. The MS2 spectrum of the 1DH intermediate species was acquired from the HPLC-purified peptide, obtained from the trypsin digest of mGsaA^B^ expressed for 6 h only. The MS2 spectrum of the ester-hydrazinolyzed 1DH intermediate was acquired similarly to the ester-hydrazinolyzed pre-albusimiditide.

### Aminopeptidase digestion of pre-albusimiditide

24 μL of the mixture containing 10 mM tricine (pH 7.0), 0.1 mM ZnSO4, and 0.5 mg/mL aminopeptidase (Sigma-Aldrich) was mixed with 3 μL of 1 mg/mL HPLC-purified pre-albusimiditide. The reaction mixture was incubated at 25 °C for 3 hours.

### mGsaA^B^ leader peptide digestion with *S. albus* J1074 culture supernatant

5 mL of GYM liquid growth medium (4 g L^-1^ glucose, 4 g L^-1^ yeast extract, 10 g L^-1^ malt extract) was inoculated with *S. albus* J1074 spores. The culture was grown for 3 days at 30 °C, shaking at 250 rpm with sterile glass beads. 1 mL of culture was centrifuged at 16,000 x *g* for 1 min. The supernatant was isolated and diluted 10-fold with sterile ultrapure water. The diluted supernatant was mixed with 10 μM mGsaA^B^ at 1:1 ratio and then incubated at room temperature for 24 hrs. The reaction was quenched and subjected to LC-MS analysis.

### In vitro isoaspartate aspartimidylation with human PIMT (*Hs*PIMT)

This procedure was adopted from our previous lihuanodin kinetics study.^39^ The reaction mixture containing 50 mM Tris-HCl (pH 7.4), 10 μM aspartimide-hydrolyzed albusimiditide, 1 μM HsPIMT, and 1 mM S-adenosyl methionine (SAM) was prepared. The reaction mixture was incubated at 37 °C for 3 hours.

## Notes

### Competing Interest Statement

The authors have declared no competing interest.

